# Selecting precise reference normal tissue samples for cancer research using a deep learning approach

**DOI:** 10.1101/336909

**Authors:** William Zeng, Benjamin S. Glicksberg, Yangyan Li, Bin Chen

## Abstract

**Background:** Normal tissue samples are often employed as a control for understanding disease mechanisms, however, collecting matched normal tissues from patients is difficult in many instances. In cancer research, for example, the open cancer resources such as TCGA and TARGET do not provide matched tissue samples for every cancer or cancer subtype. The recent GTEx project has profiled samples from healthy individuals, providing an excellent resource for this field, yet the feasibility of using GTEx samples as the reference remains unanswered.

**Methods:** We analyze RNA-Seq data processed from the same computational pipeline and systematically evaluate GTEx as a potential reference resource. We use those cancers that have adjacent normal tissues in TCGA as a benchmark for the evaluation. To correlate tumor samples and normal samples, we explore top varying genes, reduced features from principal component analysis, and encoded features from an autoencoder neural network. We first evaluate whether these methods can identify the correct tissue of origin from GTEx for a given cancer and then seek to answer whether disease expression signatures are consistent between those derived from TCGA and from GTEx.

**Results:** Among 32 TCGA cancers, 18 cancers have less than 10 matched adjacent normal tissue samples. Among three methods, autoencoder performed the best in predicting tissue of origin, with 12 of 14 cancers correctly predicted. The reason for misclassification of two cancers is that none of normal samples from GTEx correlate well with any tumor samples in these cancers. This suggests that GTEx has matched tissues for the majority cancers, but not all. While using autoencoder to select proper normal samples for disease signature creation, we found that disease signatures derived from normal samples selected via an autoencoder from GTEx are consistent with those derived from adjacent samples from TCGA in many cases. Interestingly, choosing top 50 mostly correlated samples regardless of tissue type performed reasonably well or even better in some cancers.

**Conclusions:** Our findings demonstrate that samples from GTEx can serve as reference normal samples for cancers, especially those do not have available adjacent tissue samples. A deep-learning based approach holds promise to select proper normal samples.

## Background

Comparing molecular profiles of disease tissue samples and normal tissue samples is often employed to identify a signature of the disease. The signature defined as differentially expressed genes between two groups is critical to understanding abnormal disease features and guiding therapeutic discovery [1-6]. For example, a gene expression signature created from the comparison of liver cancer tumor samples and adjacent tissue samples was used to discover anti-parasite drugs as therapeutics for liver cancer [7]. Analysis of matched tumor and normal profiles identified common transcriptional and epigenetic signals shared across cancer types [8]. Large-scale integrative analysis of cancer profiles, cellular response signatures and pharmacogenomics data suggested that such disease signatures can be widely employed for screening anti-cancer drugs [9].

However, there are many lingering issues that hinder these types of analyses. For instance, in many cancers, adjacent normal tissues are not available in these genomic databases such as The Cancer Genome Atlas (TCGA) and Therapeutically Applicable Research To Generate Effective Treatments (TARGET) (Figure 1A). As such, there is an open question on what tissue samples should be selected for these scenarios or whether creation of a proper disease signature is even possible. The recent Genotype-Tissue Expression (GTEx) project [10] has profiled samples from healthy individuals, providing an excellent resource. However, their profiles are generated from different studies and processed under different computational approaches, the feasibility of using GTEx samples as the reference remains unanswered. Moreover, given the fact there is heterogeneity within a disease, another goal is to determine a set of normal samples that are optimal for use as the reference for a group of patient samples. One approach is to choose normal samples that are similar to disease samples based on their gene expression profiles. As a substantial number of genes that are lowly expressed or not expressed at all add noises in similarity measurement, one typical alternative strategy is to utilize the top varying genes across disease samples as the features for similarity measurement. However, selecting top varying genes may ignore information of many critical genes.

**Figure 1:**
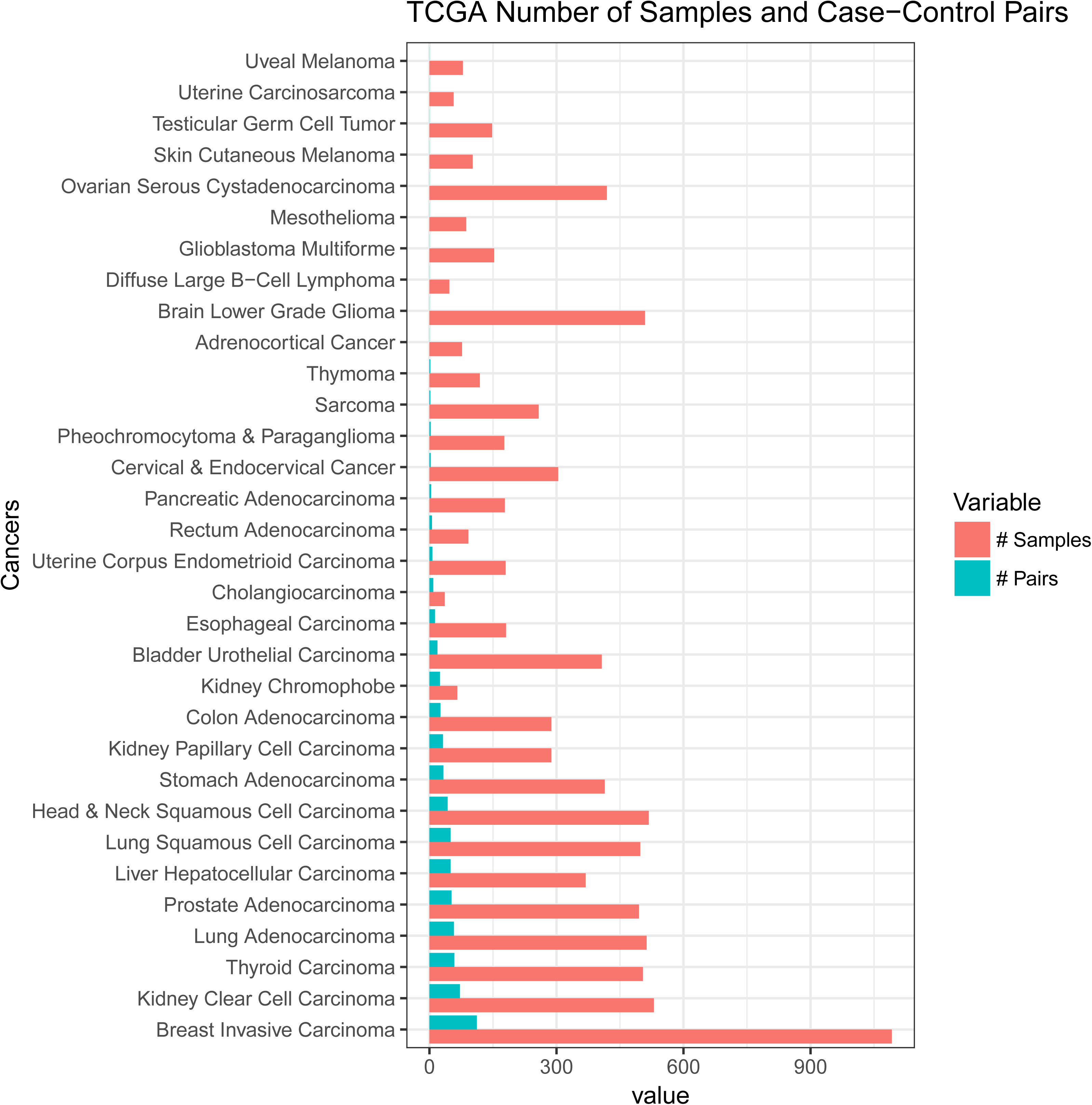
Distribution of TCGA cancer samples and pairs of case-control tissues in the dataset. Controls are adjacent tumor normal tissues.

In this work, we use the RNA-Seq data processed from the UC Santa Cruz Computational Genomics Lab’s Toil-based RNA-seq pipeline [11] and systematically evaluate GTEx as a potential reference resource (Figure 2). We use those cancers that have adjacent normal tissues in TCGA as the benchmark for the evaluation. We also explore the potential use for state-of-the-art deep learning models, specifically layers of autoencoders, to create reduced features for similarity measurement. We found that disease signatures derived from normal samples in GTEx are consistent with those derived from adjacent samples in TCGA in many cases. Our findings demonstrate that samples from GTEx can serve as reference samples for the majority of cancers, but not all. Additionally, we show promising results for utilizing deep learning strategies to select reference tissues.

**Figure 2:**
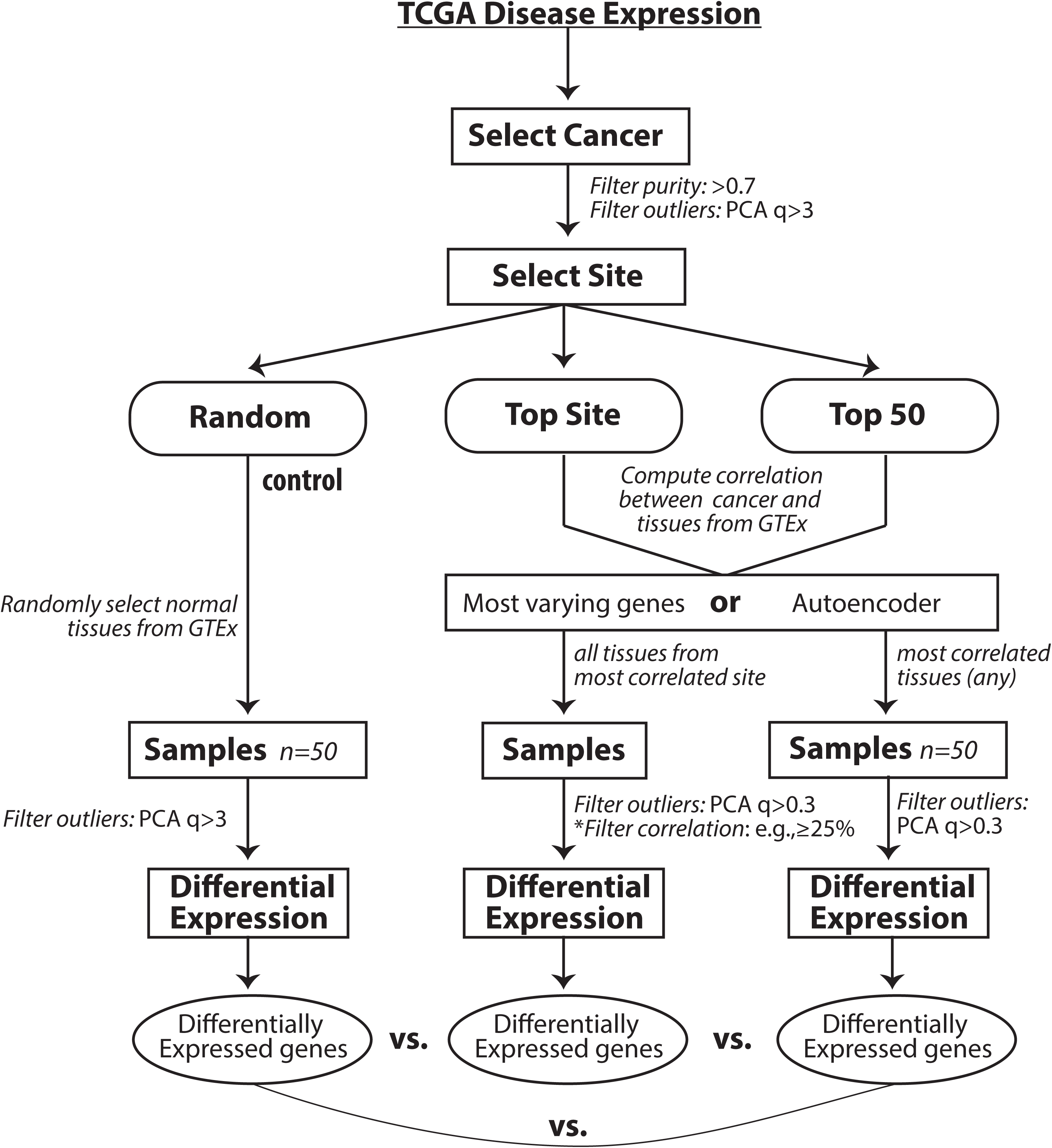
Workflow diagram.

## Methods

### Datasets

TCGA (https://cancergenome.nih.gov/) is a public repository of genomics data (e.g., gene expression) for cancer, and sometimes adjacent normal tissues. TARGET is a similar resource focused on childhood cancers. The GTEx project is a collection of gene expression data for over 7700 healthy individuals for over 50 tissues. In the current study, raw counts data and phenotype metadata for the analysis were downloaded from UCSC Xena Treehouse (https://xenabrowser.net/datapages/?cohort=TCGA%20TARGET%20GTEx) and processed into an R dataframe consisting of studies from TCGA, TARGET, and GTEx, with a total of 58,581 rows of gene expression raw counts (identified as HUGO gene symbols). Transcript abundance estimated from STAR and RSEM was used. The Treehouse raw counts data consist of 19,249 samples and, of those, a total of 19,131 tissue samples were annotated with phenotype metadata. We only used tissue samples with annotated metadata for this analysis. Of the 32 cancers, we chose cancers that have at least 10 case-control (tumor-adjacent normal) sample pairs (Figure 1).

### Workflow

In our study, we first evaluate whether our approach can identify the correct tissue site from GTEx for a given cancer (Figure 2). We then ask whether disease signatures are consistent between those derived from TCGA and from GTEx. First we selected tissues for a particular cancer in the TCGA dataset and performed quality control by filtering for tumor purity > 0.7 as determined by ESTIMATE [12]. Tissue outliers were determined by computing the principal component analysis of tissues and filtering out those with absolute z-score of the first component of greater than 3. Reference normal tissue for the tumor samples were computed using four methods:

a. Random Method: Random selection of 50 GTEx normal tissues.
b. Top Site Method: Compute correlation of GTEx tissue expression to tumor expression using top 5000 varying genes across all tumors and select all tissue samples from the top correlating tissue site. Alternatively, compute correlation using the features calculated from an autoencoder.
c. Top 50 tissues method: Compute correlation of each GTEx tissue expression to tumor expression and select the site with the tissues of the 50 highest correlation to the tumor samples.
d. Manual method: For certain tumor tissues where the computed top site did not correspond to the site of tumor. For example, if esophagus mucosa was chosen for lung adenocarcinoma, we manually selected GTEx tissue site - lung.

After the reference GTEx tissues were selected, we again removed tissues for outliers based on computed first PCA > 3. Then the tumor tissues and reference tissues were normalized using the RUVg R package library [13]. Differential expression was computed on the normalized samples. We analyzed each differential expression of the computations by comparing it with differential expressions computed from case-control set. The signature genes selected for analysis had an absolute log fold change of greater than 1 and adjusted p-value of less than 0.001.

First, we performed differential expression analysis by comparing tumor samples and normal samples using edgeR [14]. While we chose edgeR only to use, our preliminary assessment showed the conclusions hold using Limma + voom [15] or DESeq [16]. We filtered for cancers where there were at least 10 pairs of case-control samples. These were Breast Invasive Carcinoma, Kidney Clear Cell Carcinoma, Thyroid Carcinoma, Lung Adenocarcinoma, Prostate Adenocarcinoma, Liver Hepatocellular Carcinoma, Lung Squamous Cell Carcinoma, Head & Neck Squamous Cell Carcinoma, Stomach Adenocarcinoma, Kidney Papillary Cell Carcinoma, Colon Adenocarcinoma, Kidney Chromophobe, Bladder Urothelial Carcinoma, Esophageal Carcinoma. The differential expression computed from these case-control cancer samples were used as benchmark comparison to the differential expression ran against GTEx tissues as selected from the workflow (Figure 1B). To evaluate the performance we computed consistency based on the significance of overlap between signatures and correlation of fold changes of common signature genes. Further, disease expression using GTEx reference tissues were computed as in the workflow diagram. All the analyses were performed in R (version 3.4.3).

### Autoencoder

As an alternative approach to the top site methods, we evaluated the utility of an autoencoder neural network for computing correlation between cancer and reference tissue expression. Gene counts in terms of Transcripts Per Kilobase Million (TPM) from 19,260 samples were fed into an autoencoder implemented using Pytorch (v. 0.1.12_2) (http://pytorch.org/). The following parameters were used: 64 encoded features, 128 batch size, 100 epochs, 0.0002 learning rate (Figure 3A). The training took about 30 minutes using one GPU in an Amazon cloud (g3.8xlarge). Rectifying activation function, dropout and normalization were applied between layers. The loss function is defined as a mean squared error (MSE) between 60,498 elements (identified as Ensembl IDs) in the input x and output y. The functions and parameters were detailed on the PyTorch website (https://pytorch.org/docs/master/). Data were split into a training set (80%) and a test set (20%). Loss converged after 10,000 iterations (Figure 3B). The t-SNE plot of the reduced dataset shows that batch effect among three databases was minimized (Figure 3C). Similarly, we performed principal component analysis (PCA) of this datasets and chose top 64 components as the features. As the top 64 components could explain 92% variation, choosing 64 features for similarity measurement is reasonable in both autoencoder and PCA.

**Figure 3:**
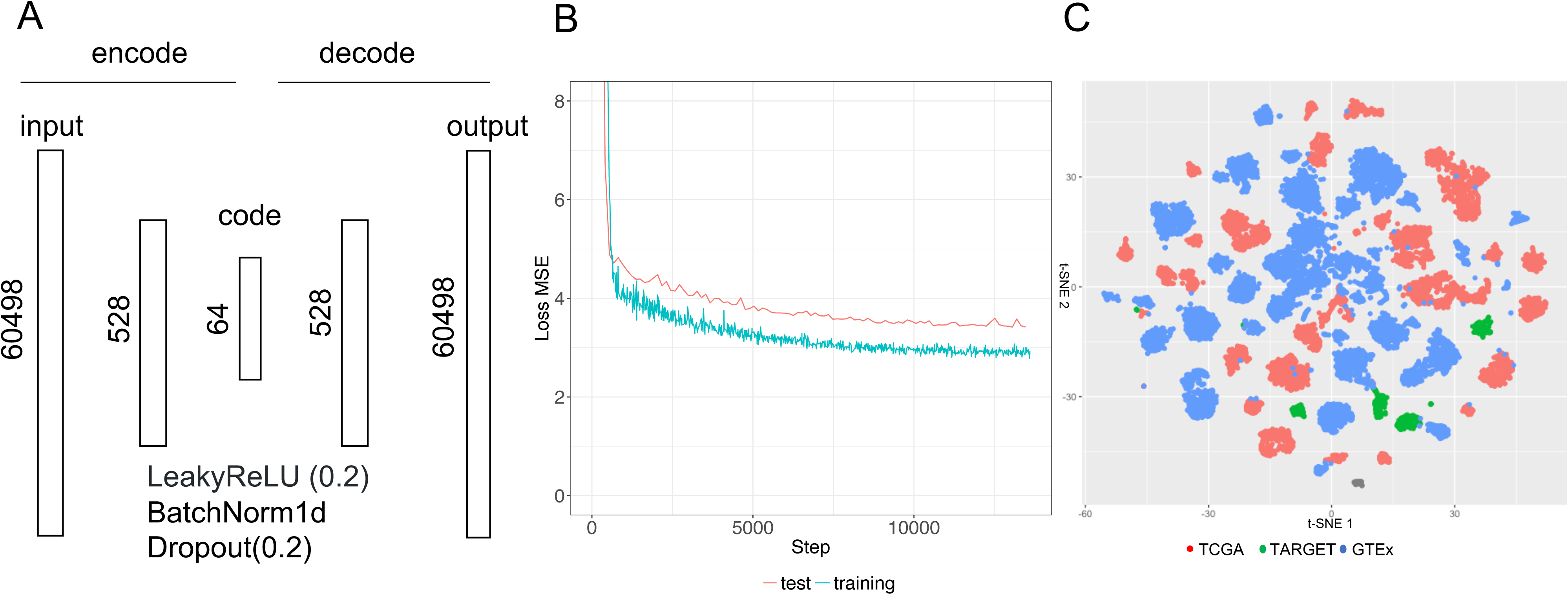
Applying an autoencoder for representing gene expression profiles. **A.** Schema and parameters. Both encoder and decoder have one layer in addition to the input/output layer. The input of encoder and the output of decoder are the expression of 60498 transcripts. The objective function is to minimize the difference between the output and input. 64 encoded features are used to represent expression profiles. Between layers, the following functions Leaky ReLU activation, batch normalization, and drop out are applied. Both network architecture and parameters can be changed. **B.** MSE loss for the training and test set. Lower MSE loss means the output is more similar to the input. **C.** t-SNE distribution of all samples using encoded features from an autoencoder. Dots were colored by data resources.

## Results

Among 32 TCGA cancers, 18 cancers have less than 10 matched adjacent normal tissue samples in the Treehouse dataset. Ten cancers do not have any matched adjacent tissue samples at all (Figure 1). Whereas, GTEx has profiles for 47 tissue sites with at least ten normal samples. This suggests the significance of exploring GTEx as a source of reference.

### Computing tissue of origin

We first asked if gene expression profiles could be used to identify tissue of origin. We indicated a site of cancer was correctly identified if the computed tissue was the site of cancer origin or a very close proximal site (potentially related site) e.g. kidney - cortex for kidney papillary carcinoma. We indicate unrelated sites as those that are further away from the cancer of origin (Figure 4). We found that using a minimal number of 100 varying genes, the correlation method can correctly identify the top tissue site for only 8 of 14 cancers. Increasing the number of varying genes to 5000 improved correct selection for 11 of 14 cancers. No further improvement on tissue selection was seen by increasing number of varying genes. The PCA, as a regular dimension reduction method, was only able to correctly identify 8 of 14 cancers, so we did not examine this method in the following analysis. The best automated method we found for reference tissue selection was via correlating autoencoder features with 12 of 14 tissues being correctly chosen.

**Figure 4:**
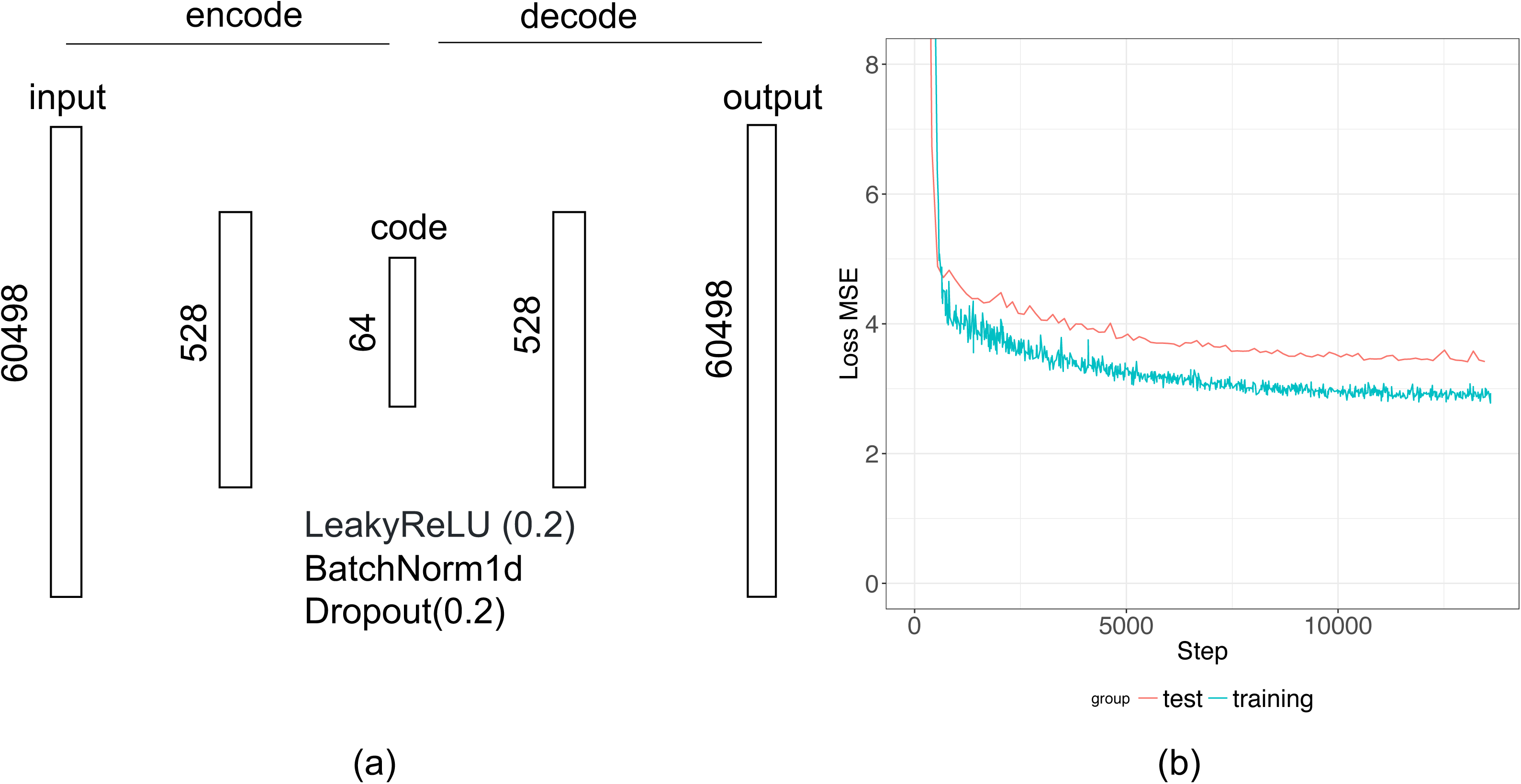
Computing tissue of origin. Top site chosen by using varying genes, PCA, and autoencoder method.

Further examination of the three misclassified cancers by varying genes methods, Bladder Urothelial Carcinoma, Lung Squamous Cell Carcinoma and Stomach Adenocarcinoma, revealed correlation values of 0.549, 0.300, and 0.858, respectively. The low correlation from the bladder and lung carcinoma may be due to substantial difference in tissue expression between the computed site, esophagus, and their expected origin site, bladder and lung. Correlation for stomach adenocarcinoma was quite high, which may be due to similarity between the computed site, ileum of the small intestine, and the stomach (Supplementary Table 1).

Squamous cell carcinomas arise from squamous cells that reside in the cavities and surfaces of blood vessels and organs. As samples in GTEx were taken from bulk tissues, this may cause the lower computed correlation between the cancer tissue and site of origin leading to erratic computational choices. Manual selection of the tissue of origin for lung squamous cell carcinoma and stomach adenocarcinoma improved the correlation from 0.549 and 0.858 to 0.883 and 0.926 respectively (Supplementary Table 1). For Bladder Urothelial Carcinoma, using the varying genes method chose esophagus - mucosa as the top site, correlation 0.549, whereas autoencoder correctly chose the bladder site, correlation. This shows that correct site choices will improve correlation.

Interestingly, Kidney Clear Cell Carcinoma, Kidney Papillary Cell Carcinoma and Kidney Chromophobe share the same tissue origin--Kidney - Cortex. This confirms that cancer can arise from different parts of one tissue and raise the question whether we should use all normal samples from one site as the reference.

### Examples of Hepatocellular Carcinoma and Bladder Urothelial Carcinoma

We use two cancers as examples for further in-depth analyses, specifically Hepatocellular Carcinoma (HCC) and Bladder Urothelial Carcinoma (BUC). In our prior results, we found that using more genes to compute the correlation generally helped to select the correct tissue site for the tumor. We ran correlation for each site using increasing number of varying genes as well as autoencoder features. We normalize the correlation of the cancer site liver (Figure 5A). We found that as the number of genes used increases all tissues will generally converge to have higher correlation with the disease tissue, this may be due to including genes of conserved regions or low expressions. Using all features from the autoencoder allows us to have much better separation of the site liver from other non-related sites of the cancer, indicating autoencoder captures the biology of disease sample more specifically (Figure 5B-C).

**Figure 5:**
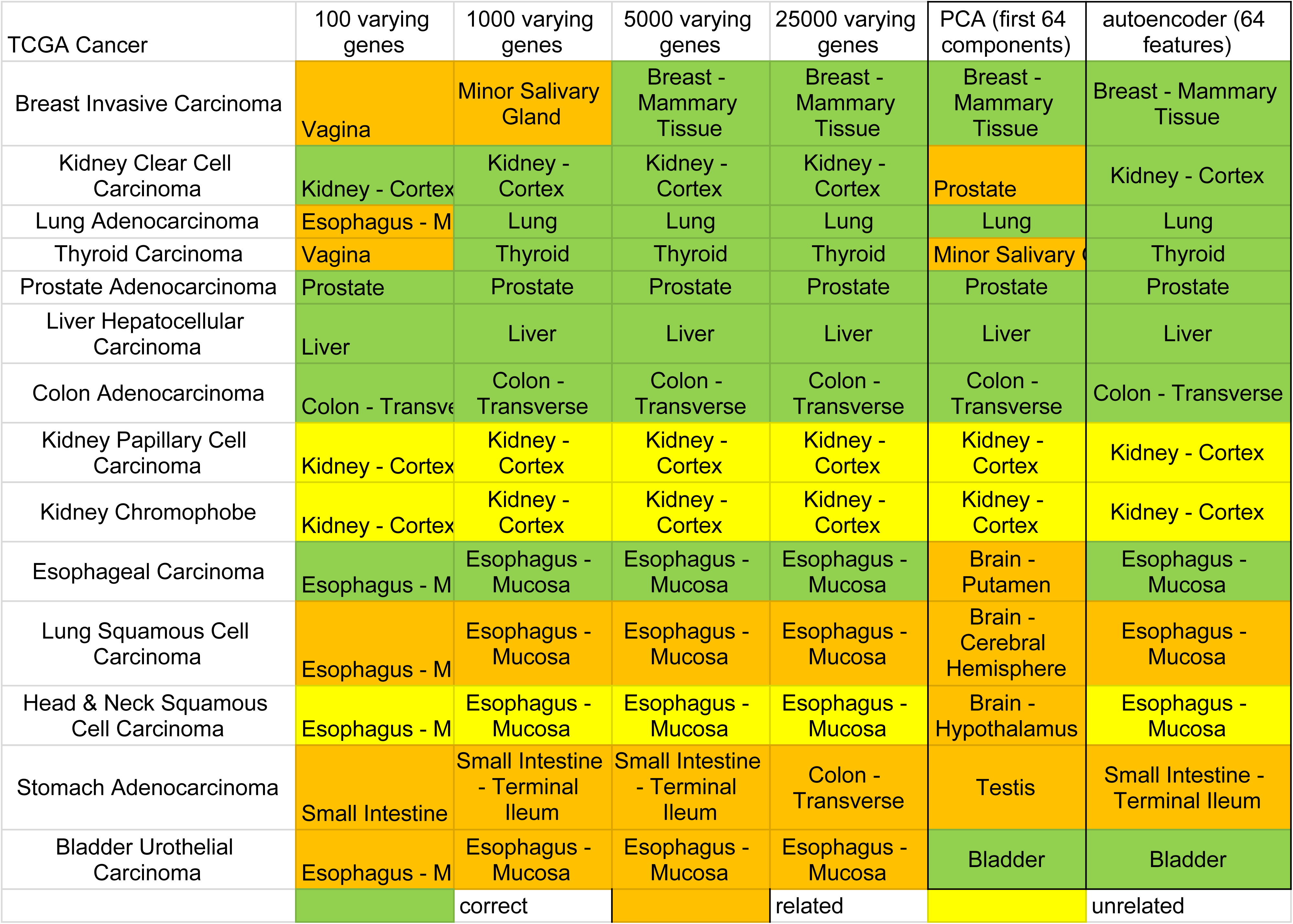
Tissue correlation between GTEx sites and HCC. **A.** Median correlation between tissue sites and cancer normalized by median liver site correlation values. **B.** Correlations between GTEx tissue sites and HCC tumor samples using top 40,000 varying genes. **C.** Correlations between GTEx tissue sites and HCC tumor samples using autoencoder features.

For BUC, however, the varying genes method was unable to determine bladder as the best site instead choosing esophagus (Figure 6A-B). Increasing varying genes from 100 to 40,000 brought down the correlation of esophagus site relative to bladder, however, it brought up correlation of other tissue sites relative to bladder (Figure 6A) similar to what we see in Figure 5A. This suggests that naively increasing varying genes does not help to distinguish tissue site selection. Meanwhile, the autoencoder method correctly predicts bladder as the top site with great separation between bladder and esophagus with greater distinction (Figure 6A, Figure 6C). Notably, the correlation in BUC is lower than that in HCC based on different similarity metrics. This suggests that cell composition in bladder tissues may be more diverse.

**Figure 6:**
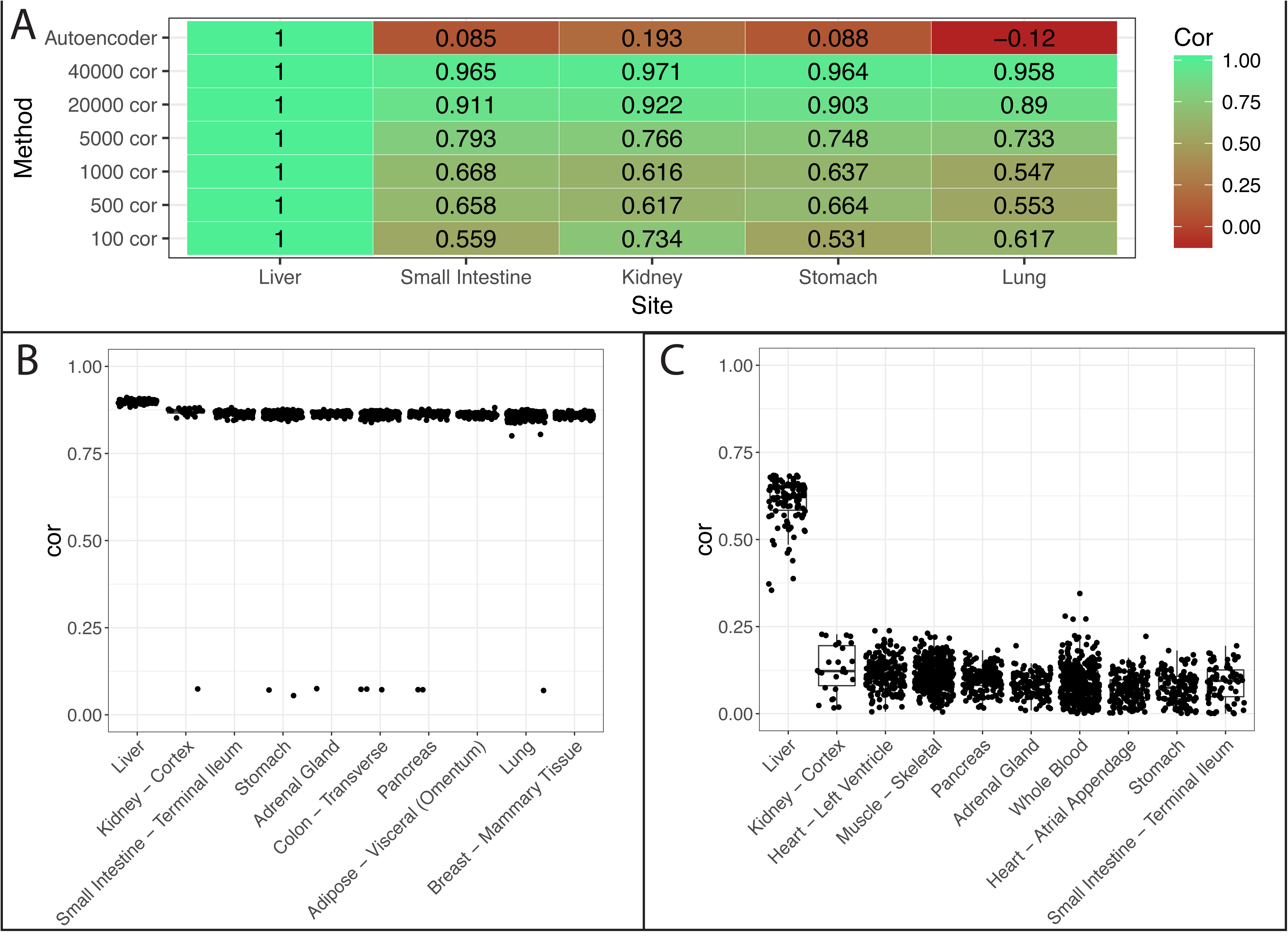
Tissue correlation between GTEx sites and BUC. **A.** Median correlation between tissue sites and cancer normalized by median bladder site correlation values. **B.** Correlations between GTEx tissue sites and BUC tumor samples using top 40,000 varying genes. **C.** Correlations between GTEx tissue sites and BUC tumor samples using autoencoder features.

### Disease Signature Comparison

As we have demonstrated that gene expression profiles can be used to identify tissue of origin, we then asked if these samples sharing the same tissue of origin from GTEx can substitute adjacent tissues from TCGA to create disease signatures. We employed three approaches to select samples (Figure 2). We evaluate consistency based on the significance of overlap between signatures and correlation of fold changes of common signature genes.

Figure 7 shows the rank-based correlation of differential expression between consensus transcripts for each cancer from TCGA using GTEx reference tissue vs. TCGA case-control samples. Using the average of three random tissue site selection as our baseline we see that our other strategies are superior. The autoencoder produced better correlations overall regardless of sample selection method.

**Figure 7:**
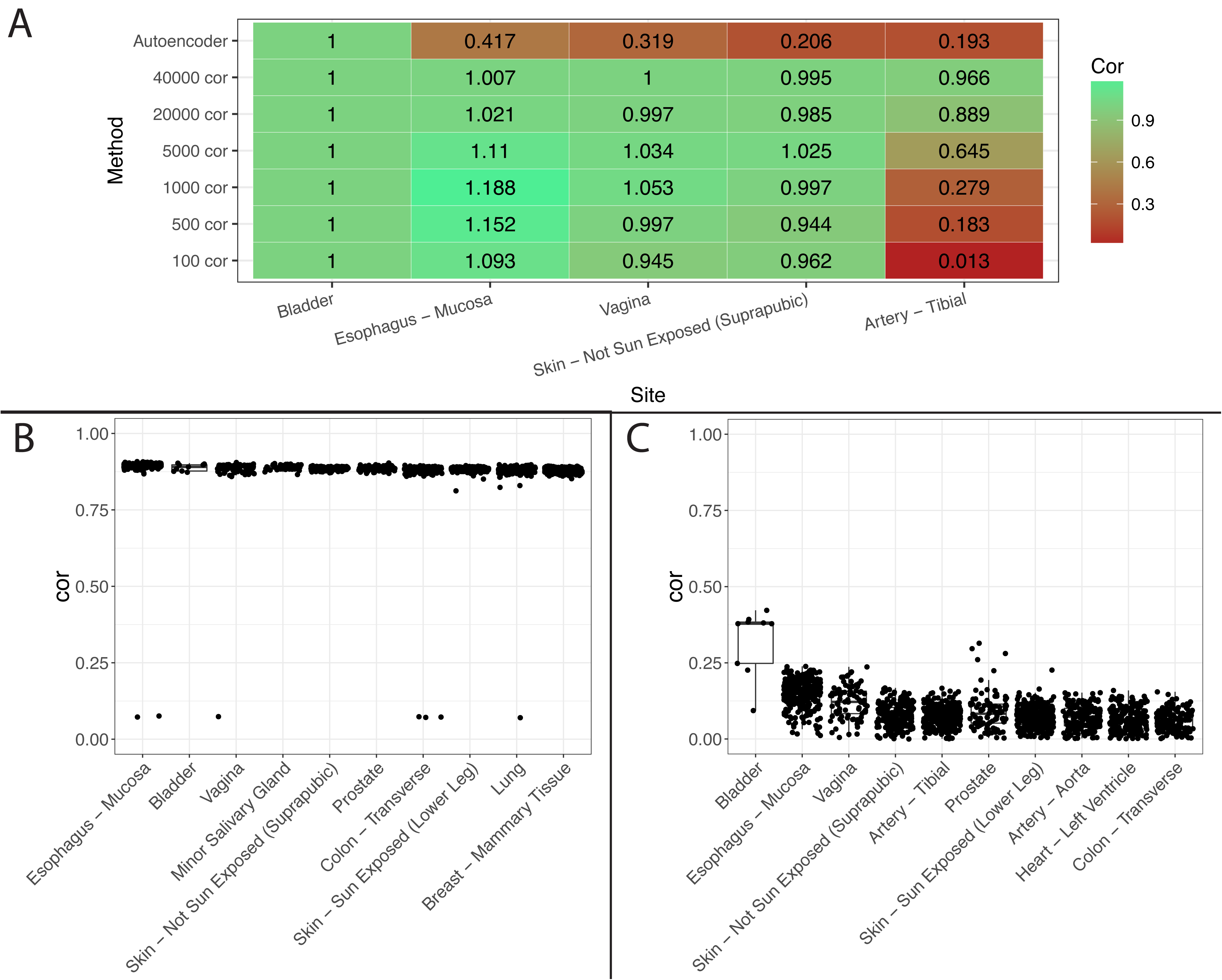
Comparing signatures from multiple methods. Auto.TopSite: choose all samples from the same tissue of origin based on autoencoder choice, Auto.25 and VarGenes.25: choose 25th percentile and above mostly correlated samples from the same tissue of origin as computed by autoencoder and varying genes, and Auto.Top50 and VarGenes.Top50: choose top 50 mostly correlated samples from all tissues as computed by autoencoder and varying genes. RandomAve: Randomly select 50 samples from all tissues. Site NA means no specified site.

For the autoencoder, it seems that choosing all samples from the same tissue of origin performs slightly better than choosing 25 percentile and above mostly correlated samples from the same tissue of origin. Interestingly, choosing top 50 mostly correlated samples from any tissue performs reasonably well or even better in some cancers, where the tissue of origin was misclassified such as the varying genes method for stomach adenocarcinoma (Supplementary Table 1). This is very significant because in many cases, where we may have no or an insufficient number of matched normal tissues, we may use normal samples from other sites. For example, in the three kidney cancers: Kidney Clear Cell Carcinoma, Kidney Papillary Cell Carcinoma and Kidney Chromophobe, our analysis suggests three cancers can share the same reference tissue sites despite the differences of origin within the kidney.

One additional question we assessed is how many normal samples are sufficient for proper disease signature-related analyses? We found that even a relatively low number of normal samples may be sufficient for calculating differential expression. For bladder urothelial cancer, for example, the autoencoder selected the bladder GTEx site which consists of only nine tissue samples (Figure 1B) for a correlation of 0.924; filtering for tissues above the 25th percentile left only seven tissue samples for a correlation of 0.926. When we used a strategy that selected more tissues, i.e. using autoencoder top 50 method, 50 sample tissues were used (9 from bladder and 41 from other top correlated sites), which produced a slight drop of correlation to 0.847. This indicates that even a relatively low number of reference tissue samples may provide a robust match.

Finally, we assessed whether it is a better strategy overall to select all samples from the same tissue site as the cancer of interest or only those that are correlated to the tumor sample. We found that the samples producing the best performance are sites where the tumor developed or a closely related site. However, when it is not possible to use such sites (e.g., when there are no available data), it is feasible to use top correlated tissues as seen from the top 50 methods. However, we found that for some cancers, even choosing top correlated sites can still produce erratic results, such as in the case of lung squamous cell cancer. In this case, the correlations for all non-random methods were between 0.1 - 0.3 was not even able to beat the random tissue selection (Supplementary Table 1). Along these lines, we evaluated differential expression similarity using samples from a different origin than the cancer of interest. For example, in two kidney cancers, Kidney Papillary Carcinoma and Kidney Chromophobe the kidney cortex were computed as the top site, for Head and Neck carcinoma the esophagus-mucosa was the top site. Their high correlation with case-control >0.8 indicates that choosing sites at different origin but proximal to the cancer will provide good disease signature (Supplementary Table 1).

### Assign normal tissues for cancers with low case-control pairs

Since there were 18 cancers with insufficient number of adjacent normal tissues, we use our computational approach to assign a primary site for each. Of the 18 cancers, the autoencoder method was able to determine 10 correct sites, whereas using the top 5000 varying genes only produced 4 correct sites (Figure 8). This suggests an autoencoder can select proper samples to create disease signatures for those cancers.

**Figure 8:**
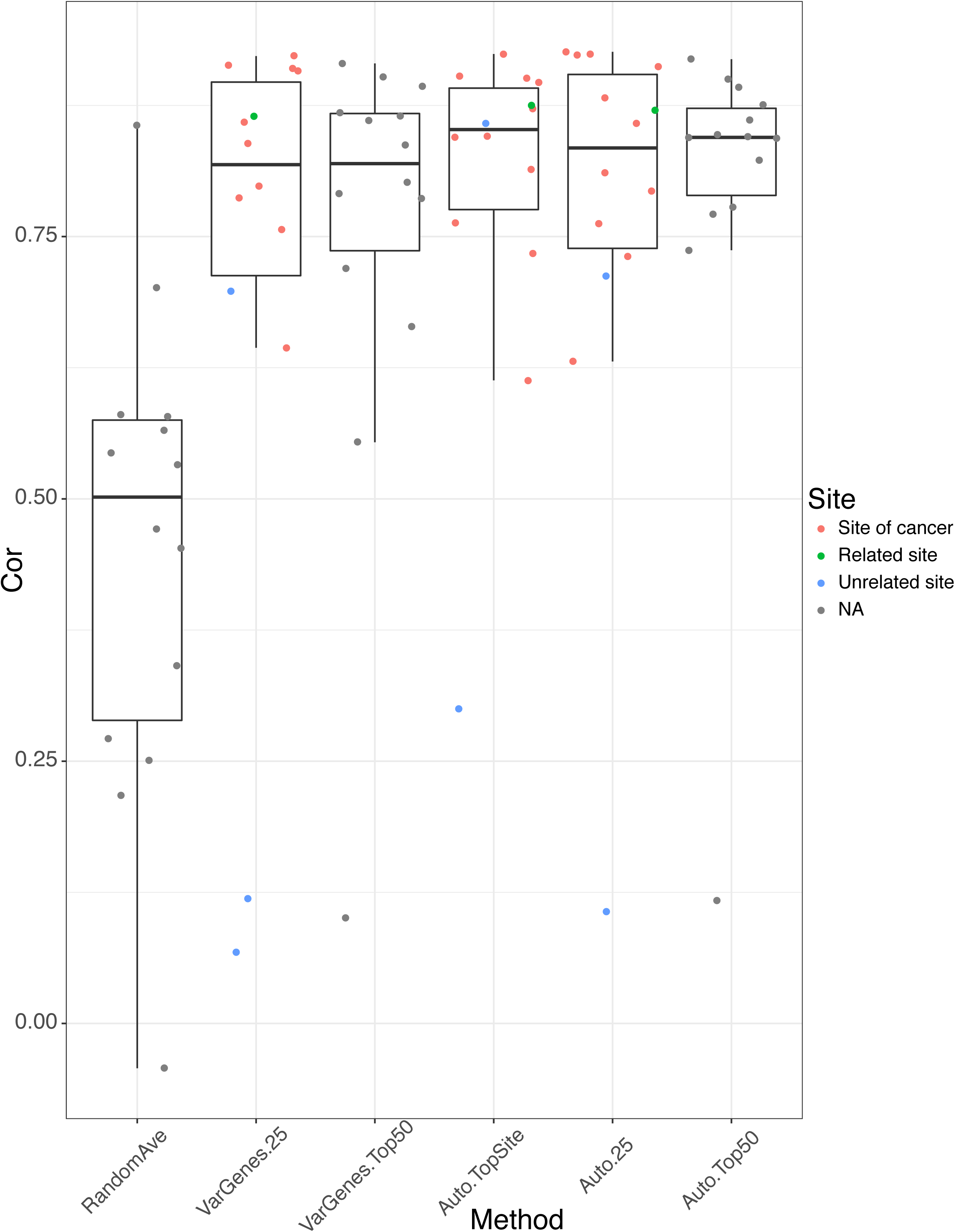
Putative site computed for cancers with low case-control pairs using 5,000 varying genes and autoencoder features.

## Conclusions

In the current study, we evaluated the nuances of proper reference tissue selection for disease signature-related analyses. Furthermore, we assessed the benefit of using state-of-the-art methodologies, namely deep learning via an autoencoder strategy, to enhance performance of identifying ideal reference tissues for cancers of interest.

The findings from our study will significantly enhance probing disease biology through gene signatures. As the cost of sequencing is rapidly decreasing, it becomes very common to profile disease samples of interest, however, collecting matched normal tissues from patients is difficult in many instances. Our analysis confirms that GTEx, the largest cohort of normal samples, can serve as a source of reference normal tissues in cancer research. In the current study, we chose to focus on cancer because we have plenty of adjacent tissues that can be used as a benchmark. We expect that the methods and findings from our study can be extended for cancer subtypes or other non-cancer research as well. However, a few caveats have to be considered. First, all RNA-Seq data have to be processed in the same pipeline in order to mitigate batch effects. Second, some disease samples may have no relevant normal tissue samples because of diverse cellular composition. This limitation may be addressed by using cellular decomposition techniques or single cell data.

Based on the success of this study, we have some future works that we are exploring. Although we show the potential of using autoencoder for feature selection, we have not fully optimized the model for tissue selection. In our exploratory studies, we found that encoded features are very sensitive to network architecture and parameters, although it does not affect the results in the computation of tissue of origin. For example, when we changed learning rate from 0.0002 to 0.005, batch size from 128 to 64, dropout rate from 0.2 to 0.1, LeakyReLU negative slope from 0.2 to 0.1, respectively, the average correlation between the new features and the default features changed to 0.219, 0.069, 0.354, and 0.219 (Supplementary Table 2). Interestingly, while a new layer was added into the network, the average correlation even decreased to −0.01. However, while using new features to compute tissue of origin, we observed that all new features could clearly separate the first top site and the second top site. For example, in liver and bladder cancers, liver and bladder are predicted as the top site respectively, and the correlation with the top site is much higher than that with the second site (Supplementary Table 2). Surprisingly, when the feature size was reduced from 64 and 32 or two new layers were added, the top site of bladder cancer was incorrectly predicted. In short, given the complexity of neural networks, additional effort should be made to optimize the model, nevertheless, we indeed demonstrate the superiority of deep learning models in this work.

Furthermore, in addition to using gene expression as features, we will explore adding other cancer specific features including presence of mutations and copy number variation. The autoencoder strategy would be able to manage such diverse feature types. We also plan to determine whether changing the order of workflow, such as removing outliers first, might improve this analysis. In addition, as adjacent cancer normal tissues are sampled near the cancer site, some of these tissues may contain cancer cells and thus have some expression of cancer[17], which may require further investigation. We will further explore our approach to study pediatric cancers (available in TARGET), where adjacent normal tissues are even more scarce.

## List of abbreviations

PCA =: Principal Component Analysis
TCGA =: The Cancer Genome Atlas
TARGET =: Therapeutically Applicable Research To Generate Effective Treatments
GTEx =: Genotype-Tissue Expression project
RNA-seq =: RNA sequencing
HCC =: Hepatocellular Carcinoma
BUC =: Bladder Urothelial Carcinoma

## Declarations

### Authors’ contributions

WZ and BC conceived the study. WZ performed the majority of analysis with the input from BG and BC. YL and BC implemented the deep learning infrastructure. WZ, BG, and BC wrote the manuscript with the input from YL. BC supervised the study.

**Authors’ Information**

## Availability of data and materials

Data and code will be available upon request.

## Competing interests

No conflicts of interests are declared.

## Declaration

The research is supported by R21 TR001743, U24DK116214 and K01 ES028047 (to BC). The content is solely the responsibility of the authors and does not necessarily represent the official views of the National Institutes of Health.

## Tables

None

## Additional Files

Supplemental Table 1: Consensus sequence and gene rank correlation with case-control pairs using different methods. Differentially expressed genes were selected using adjusted p < 0.001 and absolute log fold change > 1. Consensus sequences are defined as overlapping differential expression sequences with same directionality in log fold change. Rank correlation is the Spearman’s rank correlation of differential expression (fold change) between the consensus sequences computed from multiple methods (see workflow) and case-control pairs. Unless otherwise stated all rank correlation have p values < 0.01.

Supplemental Table 2: Autoencoder architecture and parameter evaluation. New encoded features were used to compute tissue of origin. A better separation between the first top site and the second top site indicates a better model.

---

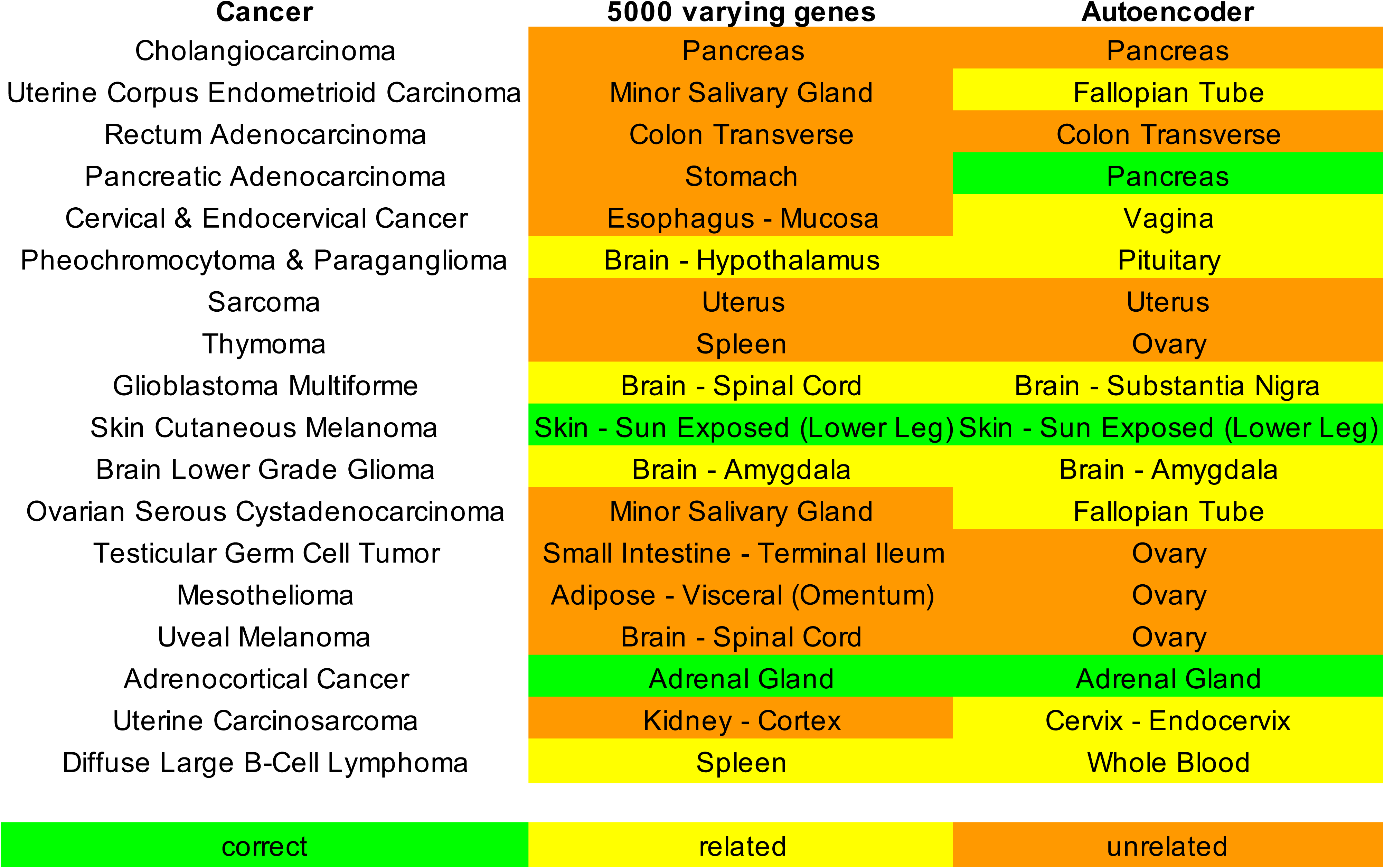

**Figure 10:**
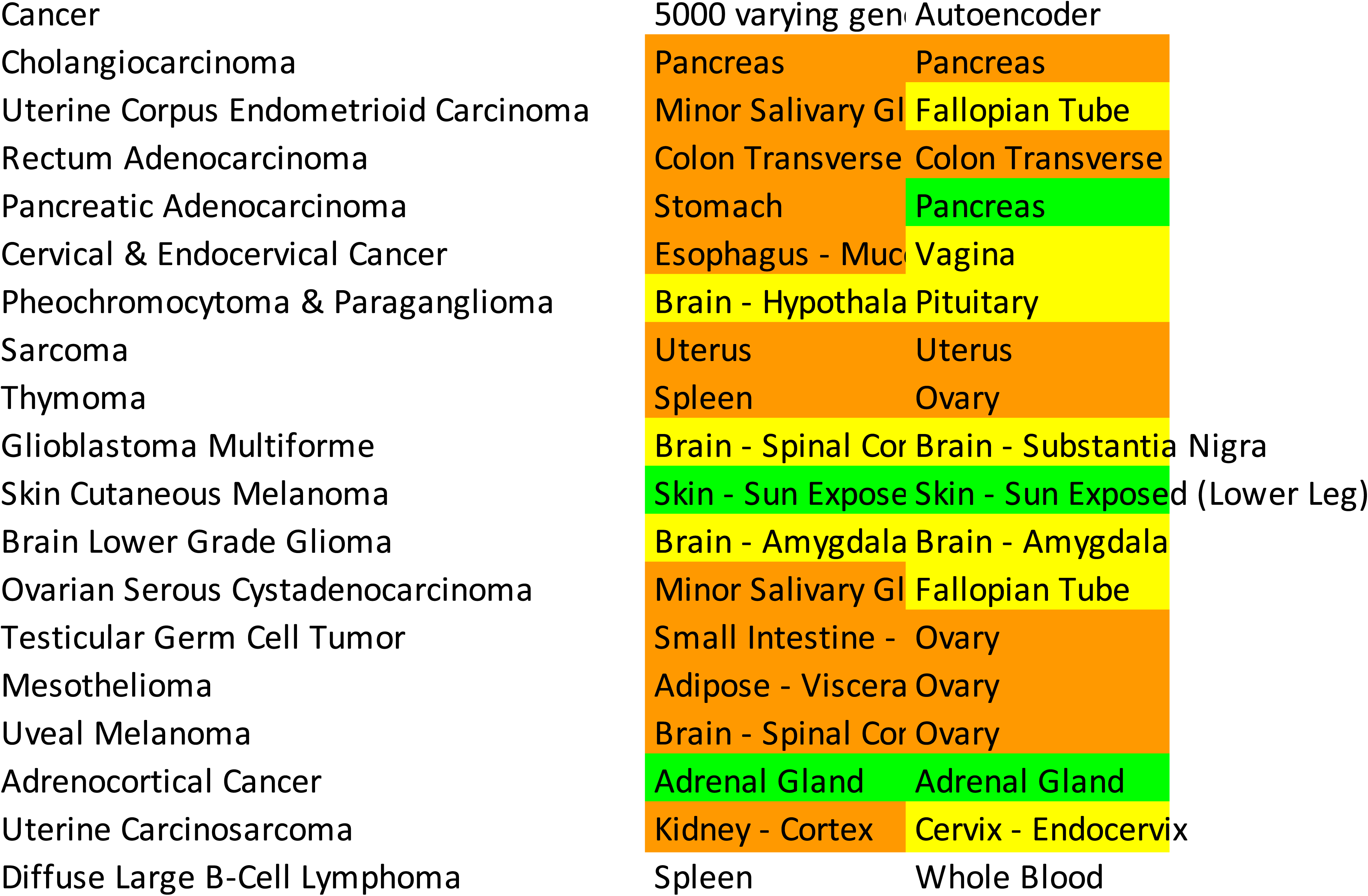

